# Parental care modifies the role of early-life size and growth in shaping future physiology

**DOI:** 10.1101/2024.09.27.615193

**Authors:** Zachary M. Laubach, Sage A. Madden, Aleea Pardue, Rebecca J. Safran

## Abstract

Size and growth early in life are associated with physiological development and these traits influence fitness. Life history theory predicts that the relationship between traits reflect constraints involving allocation and acquisition of resources. Using longitudinal data from 113 wild nestling barn swallows (*Hirundo rustica erythrogaster*), we first characterized developmental changes in glucose metabolism, a physiological trait involved in energy mobilization and response to stress. Next, we used these data to test hypotheses from life history theory about allocation and acquisition of resources based on associations of nestling size and growth with glucose physiology and assessed whether these relationships are modified by parental care. We found that larger nestlings had higher baseline blood glucose and larger magnitude of change in glucose in response to a stressor. Further, this relationship was most pronounced among birds in nests that received the lowest amount of parental care. Given that glucose metabolism fuels activity and is critical in the vertebrate stress response, these results suggest that physiological constraints may contribute to the early-life disadvantage of being smaller, especially in the context of lower parental care. While these findings are inconsistent with a trade-off involving differential allocation of resources between life history traits, they align with the differential acquisition hypothesis.

## INTRODUCTION

Larger size and faster growth early in life are hypothesized to improve fitness by increasing survival to adulthood and thereafter, facilitating reproduction (Stearns and Koella 1986; Stearns 1992). Empirical evidence from diverse taxa, including birds and mammals, supports this hypothesis (Lindström 1999; Ronget et al. 2018). In birds, juvenile size and patterns of early-life growth correlate with ontogenetic development of physiological mechanisms (Cuervo et al. 2011; Lill et al. 2013), which, in turn, affect performance, *e.g.*, flight ability at fledging (Cornell et al. 2017) and survival (Bowers et al. 2014; Brown et al. 2021). Traits that affect fitness, including structural growth and physiological function, require adequate resources such as nutrition during early life (Stearns 1989). When resources are limited, the associations between life history traits are expected to follow one of two general patterns that reflect differences in resource allocation and acquisition (Van Noordwijk and De Jong 1986). First, if faster growth corresponds with reduced physiological capacity, then this negative association indicates a life history trade-off (Allen et al. 2022) in which individuals differentially allocate limited resources between competing traits such that an investment in one trait results in neglect of another (Stearns 1989). Second and alternatively, larger size may be associated with more robust physiological capacity (Westneat et al. 2004; Forsman et al. 2010). In this scenario, it is hypothesized that resources are similarly allocated between traits, but that individuals differ in the amount of resources they acquire, resulting in positive trait covariation (Van Noordwijk and De Jong 1986). Theory and empirical evidence support both negative and positive associations between structural and physiological traits, suggesting that individuals may face constraints involving differential allocation and acquisition of resources.

In altricial birds, fledging is a key life history event since it requires sufficient growth and development of traits necessary for flight (Cornell et al. 2017) - a transitional milestone marked by high mortality (Maness and Anderson 2013; Cox et al. 2014; Martin et al. 2018; Jones and Ward 2020). As such, the life stage(s) prior to fledging is likely a sensitive period during which investment of energy into structural components, including wing length – a key indicator of flight ability (Cornell et al. 2017) – and physiological development are crucial to survival.

Glucose metabolism is integral to juvenile development and the ability to fledge, in part because glucose fuels activity (Braun and Sweazea 2008) and because glucose is a primary downstream product of the hypothalamic pituitary adrenal (HPA) axis response to acute stressors (Kuo et al. 2015; Vágási et al. 2020). In birds, baseline blood glucose concentrations are maintained at relatively high levels to support essential functions like brain metabolism (Scanes and Braun 2013; Sweazea 2022). Glucose metabolism in response to a stressor is also an important indicator of vertebrate stress physiology. Immediately following exposure to a stressor, catecholamines mobilize energy stores, causing blood glucose levels to increase. Glucose levels remain elevated, often for several hours, via glucocorticoids inhibiting cellular uptake and glucose storage (Sapolsky et al. 2000). This coordinated action of the HPA axis is one mechanism through by which organisms adaptively allocate energy to respond to and potentially recover from acute stressors (McEwen and Wingfield 2003; Romero et al. 2009; Romero and Beattie 2022).

Avian glucose metabolism at baseline and in response to an acute stressor represent two facets of physiological mechanisms that are sensitive to an individual’s condition and age (Scanes and Braun 2013; Vágási et al. 2020). Specifically, fasting state blood glucose provide information on an individual’s capacity to fuel essential metabolic functions whereas stress-induced glucose levels shed light on an individual’s ability to respond to and recover from stressors (Remage-Healey and Romero 2000; Romero and Beattie 2022). Characterizing changes in baseline glucose over the course of development and in response to a stressor can provide proximate explanations for this physiological trait. Such efforts can also enable investigating how basic physiological mechanisms, like glucose metabolism, are a related to life history strategies (Ricklefs and Wikelski 2002), thus generating insight into ultimate explanations for the relationships between life history traits.

We identify several key gaps in current knowledge regarding the relationships between nestling size and growth with physiological development early in life, and how parental care affects those relationships. First, while structural and physiological trait associations are likely to change over the course of development, most published literature focuses on these associations at a single point in time. Repeated assessments of traits across early life are needed for an ontogenetic understanding of individual trade-offs and constraints that influence structural and physiological development (Allen et al. 2022). Second, associations between offspring size and physiology occur in broader social contexts that have historically been overlooked in studies linking early-life size and growth to physiology. Social factors, like parental care and number of siblings, can modify the association between structural and physiological development by influencing nestlings’ acquisition of resources (Ricklefs et al. 1998). In birds, a lower per nestling provisioning rate and nutritional restriction are associated with smaller size (Boag 1987; Searcy et al. 2004; Martin 2015; Sofaer et al. 2018) and poorer physiological condition (Burness et al. 2000; Pravosudov and Kitaysky 2006; Bańbura et al. 2008). Still, less is known about how parental care influences the association between structural and physiological traits during early development, and whether empirical patterns align with predictions from life history theory.

We used longitudinal data from 113 wild nestling barn swallows (*Hirundo rustica erythrogaster)* from 31 nests to investigate the relationship between nestling size/growth and early-life physiology. We focused on wing length as a morphological trait that correlates with body size and blood glucose levels as an indicator of physiological development as higher levels of circulating glucose occurs with successful energy mobilization and adequacy of the stress response (fasting blood glucose levels; **Figure 1**). Since studies on glucose physiology in wild nestlings are limited, we first asked, does nestling blood glucose increase in response to an acute stressor, and are individual glucose measurements consistent across development? To answer these questions, we characterized nestlings’ fasting blood glucose response to an acute stressor (**Figure 1, Step 1**) and used repeated samples across nestling development (∼8 days old and ∼12 days old) to assess the intraindividual repeatability of circulating glucose measures (**Figure 1, Step 2**).

**Figure 1.**
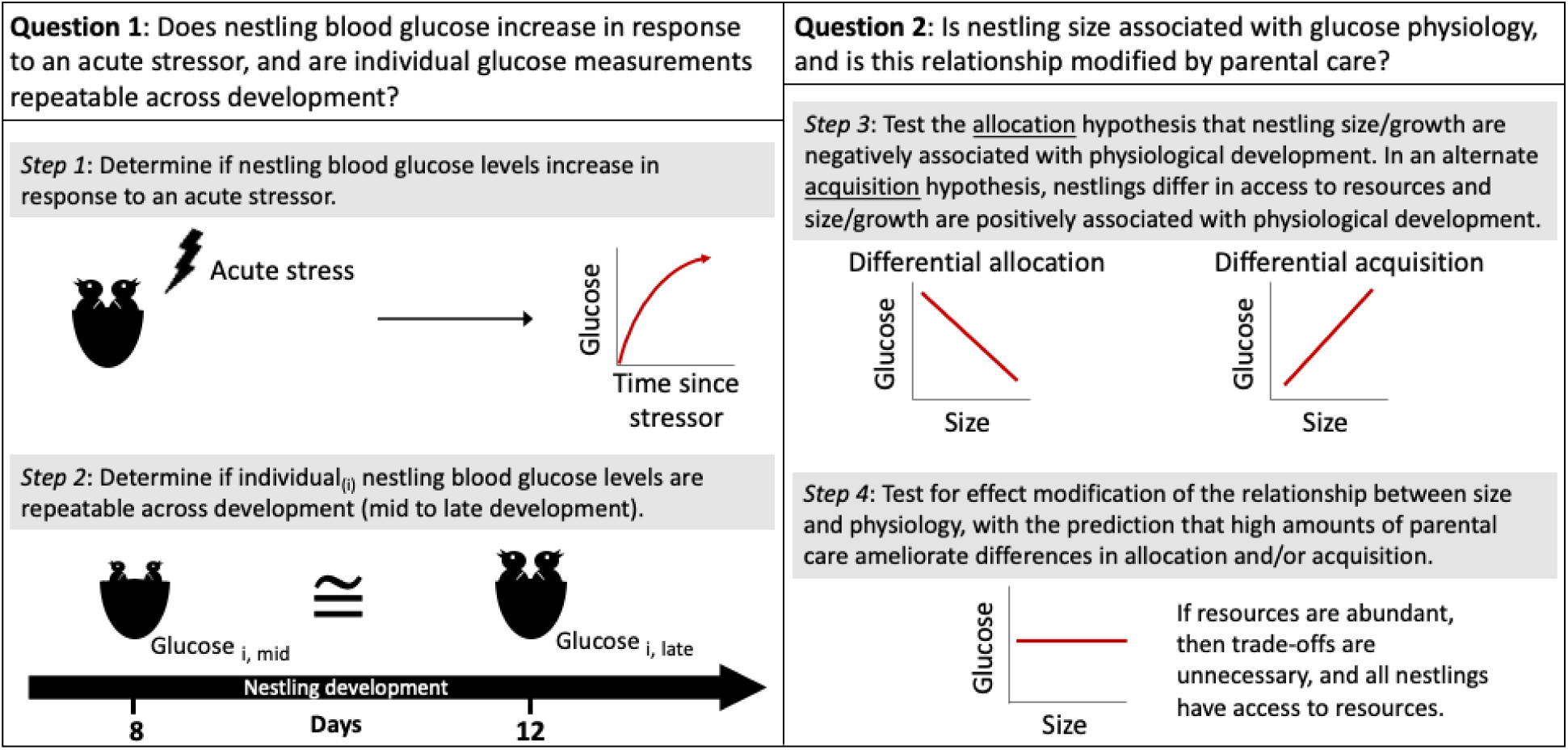
A conceptual diagram of the research questions and possible outcomes from the analytical steps taken to characterize fasting blood glucose levels and to test hypotheses about the relationships between size, physiology, and parental care during early development. Predicted relationships from Step 3 are based on a simplified version of Figure 1 by van Noordwijk and de Jong. 1986. *Am. Nat*.

Next, we asked is nestling size and/or growth associated with current and future blood glucose levels, and is this relationship modified by parental care? Here, we investigated associations between nestling size/growth and glucose levels (**Figure 1, Step 3**). Drawing on life history theory, we hypothesized that larger size and greater growth would be associated with limited glucose metabolism, including lack of an increase in circulating blood glucose levels in response to an acute stressor, if differential allocation of resources between structure and physiology required a trade-off. Alternatively, smaller nestlings might be limited in their ability to regulate their glucose metabolism. This alternative pattern would be expected if differences in resource allocation are small relative to interindividual differences in resource acquisition. Finally, we investigated nest level measures of parental care as an effect modifier to the relationship between size/growth and glucose (**Figure 1, Step 4**).

For Steps 1-4 in this paper, we employ distinct data analysis tasks of description (i.e., characterization of the central tendency and spread of continuous variables; assessing the % of subjects across categories of discrete variables), association (*i.e.*, establishing a relationship between X and Y), and causal inference (*i.e*., testing whether X causes Y) (Hernán et al. 2019; Laubach et al. 2021; Tredennick et al. 2021). For Steps 1 and 2, we aimed to characterize the glucose response mechanism and glucose repeatability using a combination of description and association. The descriptive analyses provide population and subgroup averages and corresponding measures of biological variation to summarize the data. The association analysis leverages basic knowledge on temporal ordering of variables to observe relationships between two variables. These prior steps then inform Steps 3 and 4, which involve use of expert *a priori* knowledge on the potential causal relationship between nestling morphological traits and glucose over time. For Step 3, we identified and controlled for potential confounding variables that, if not accounted for, may lead to biased estimates. For Step 4, we assessed for effect modification by parental care. This analysis aligns with our hypothesis that better quality or quantity of parental care may buffer unfavorable effects of small size or sub-optimal early-life growth on physiology (Laubach et al. 2022). Specifically, we hypothesized that high amounts of parental care would ameliorate the effect of size/growth on glucose metabolism following the prediction that abundant resources limit the need for trade-offs and reduce resource access inequity among nestlings.

## METHODS

### Study population

We collected data on demography, morphometry, physiology, and behavior from migratory barn swallows during their 2021 summer breeding season at 7 established breeding sites (farms) in Boulder County, Colorado. These sites are monitored as part of a long-term study of barn swallows and are occupied by between 1 to 50 breeding pairs. Females typically lay clutches of 3-5 eggs that, following incubation, hatch asynchronously over approximately 1-2 days. This asynchrony results in a size hierarchy that persists over the course of development (Saino et al. 2001). Hatching order is not associated with offspring sex in this species, and there are no differences between female and male nestlings when comparing multiple morphological or physiological (serological and immunological) traits (Saino et al. 2002). Finally, from the time eggs are laid until nestlings fledge (∼18 days post hatching), socially paired female and male parents provide care, which includes incubation, brooding, nest sanitation, and food provisioning (Møller 1994; Brown and Browon 2020).

### Sampling design and data collection

For this study, we monitored 113 nestlings from first clutches from 31 nests during the months of May – July 2021. We collected data following a prospective sampling design (**Figure 2**). Nests were monitored twice per week and nestlings’ ages were estimated upon hatching based on the emergence and visibility of feather tracts, whether the nestling was found wet, and the ability of the nestling to raise its head (Fernaz et al. 2012). Each nest was assigned an age, beginning with day 0 on the date when the first nestling hatched. Nestling morphometry, nestling ectoparasites counts, nest temperature, and parental care data were collected at early development (day 3-4), mid-development (day 8-9), and late development (day 11-13). Near nest temperatures were collected from Govee ® thermometers hung in small mesh bags adjacent (between 10 – 28 cm) to each active nest. Here, we focused on the median ambient temperature recorded during the parental care observation since temperature may affect nest activity including parental care behaviors. Nestlings were banded with uniquely numbered aluminum leg bands at mid-development. Blood samples were collected at mid- and late development.

**Figure 2.**
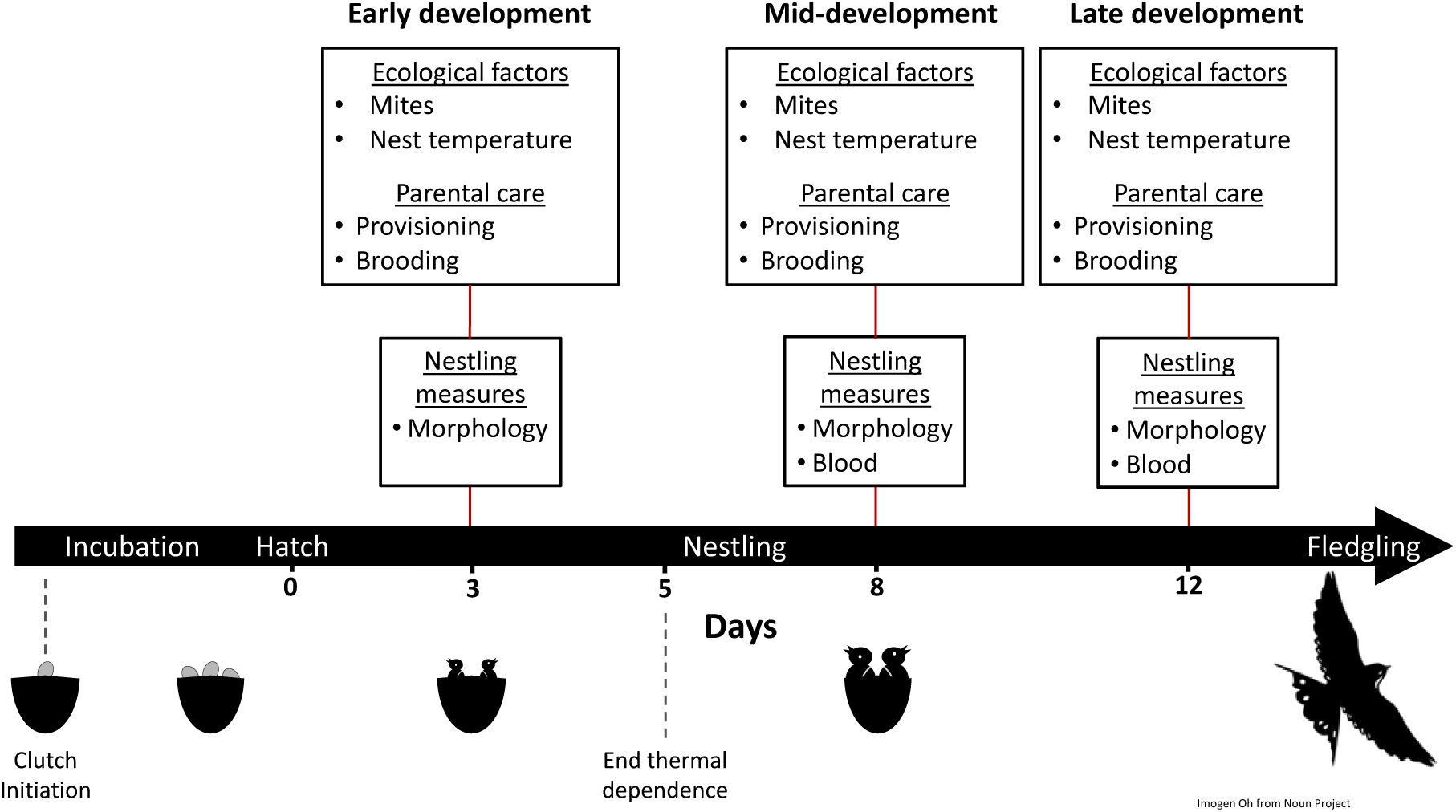
An overview of the sampling design indicating the developmental state and the types of data that were collected from wild barn swallows.

### Assessment of independent variables: nestling morphometric measures

We collected standard morphometric measurements from each nestling. We measured mass to the nearest 100^th^ of a gram using a digital scale, and right-wing length to the nearest half millimeter as the distance from the carpal joint to the distal end of the second phalanx or primary wing feather, whichever was longer. We also created a categorical variable of relative nestling size, where individual nestlings were categorized as ‘small’ if their right-wing length was below the within nest average right-wing length, and ‘large’ if above the average. We set small relative size as the reference group for all downstream analyses. Ectoparasites, which primarily included blood mites, were counted by visually inspecting nestlings. Due to low counts, we summarized these measurements as a binary variable indicating presence or absence of ectoparasites.

### Assessment of dependent variables: blood glucose levels

In this study, we were interested in two assessments of blood glucose: baseline levels and change in glucose levels before vs. after an acute stressor. We interpreted higher fasting baseline glucose and a larger magnitude of change in glucose in response to a stressor as favorable outcomes, reflecting greater energetic reserves and the ability to mount an adequate stress response, respectively.

To assess baseline glucose levels, we measured fasting blood glucose levels for each nestling at mid- and late development. To estimate change in glucose levels between baseline and an acute stressor (*i.e.,* stress response glucose), we subjected each nestling to an acute stressor which entailed removing nestlings from their nest and handling them while measuring morphology. Both baseline and stress response blood glucose were measured between 3:25 – 5:56 am, while nestlings are presumed to be fasting since barn swallows are aerial insectivores that feed in daylight. Corticosterone begins to increase in passerine birds in response to an acute stressor within approximately 3 minutes (Romero and Reed 2005), and considering that circulating glucocorticoids increase blood glucose, we attempted to measure at near baseline glucose levels. For the baseline blood glucose levels, the average (range) sampling time, which is the time between extracting the first nestling from the nest and the time at blood sampling, was 2 minutes 46 seconds (52 seconds – 4 minutes 58 seconds), except for 6 samples that were taken between 5 and 10 minutes. The average (range) sampling time for the stress response blood glucose measurements was 19 minutes and 26 seconds (16 minutes – 27 minutes 16 seconds), which is a long enough time following a stressor to capture an increase in blood glucose levels (Remage-Healey and Romero 2000; Montoya et al. 2020), while minimizing the time nestling were removed from their nests. We collected a few drops of blood from the brachial vein and glucose levels were measured with a FreeStyle Lite portable glucose meter (Abbott Diabetes Care, Alameda, CA, USA). Glucose measurements from the FreeStyle meter have been validated in closely related species (Taff et al. 2021, 2022).

### Assessment of the effect modifier: parental care

We measured parental care behavior by conducting ∼1 hour long focal observations that began between 5:45 – 6:53 am in the morning, except for two sessions that began at 7:28 and 8:00 am. The start of each focal observation proceeded collecting morphological and physiological data from nestlings and a subsequent ∼15-minute habituation period (**Figure 2)**. The median (range) duration of time between extracting nestlings from the nest and initiating the focal observation was 77 (23-219) minutes. Once parents were no longer visibly agitated or alarm calling, trained observers concealed in a blind recorded counts and durations of behaviors from both female and male social parents using the Animal Behavior Pro application, version 1.6 (Newton-Fisher 2021). In other words, behavioral observations and data collection began only after adult birds did not appear disturbed by our presence. It is worth noting, given that these birds are used to humans in the barns the return to normal behavior often happened quickly. When an observer could not be present, we recorded nests using GoPro® cameras and videos were scored by the same observers who conducted the in person focal observations. When using a camera, we set up the camera mounts the day prior to recording so that the birds could habituate. We have previously shown that parental care behaviors scored by different observers and between in-person versus camera recordings are consistent (Madden et al. 2022). A full ethogram describing each behavior is included in the supplemental information (**Supplemental Table 1**). Parents were identified via colored leg bands and unique morphological traits, including tail streamer lengths and breast plumage coloration.

We created an index of parental care by summarizing feeding counts from both parents across the three stages of nestling development (**Figure 2)**. Here, we focused on feeding behaviors because this provides a measure of resource availability at each nest, which according to life history theory is predicted to influence the relationship between traits (Laskowski et al. 2021). More specifically, we used mixed effects models, in which total feeding counts by both parents measured at early, mid-, and late development were included as a repeated measures outcome and nest ID was a random intercept. We compared model fit for multiple distributions and chose a negative binomial for our final model (**Supplemental Table 2**). In this model, we included an offset for total observation time and adjusted for several variables that could differentially affect parental care behaviors. These covariates included the number of days since the first nestling hatched, the number of nestlings in a nest, the median temperature near the nest during the observation trial, and the duration of time between extracting nestlings from the nest and the start of the focal data collection. From this model, we extracted the best linear unbiased predictors (BLUPs), which provide an inter-nest ranking of parental care based on nest-level average feeding rates summarized across development (**Figure S1**, in supplemental materials). In addition to controlling for environmental conditions and nest-level variation that may influence parental feeding rates, using a mixed model to summarize parental care was efficient since it allowed for missing data via shrinkage of nest-level estimates towards the overall sample mean. Further, because parental care is analyzed as an effect modifier, we created a two-level categorical variable by classifying the BLUPs as either low or high parental care. We note that the approach of carrying forward BLUPs from one model to another has received criticism for its failure to propagate uncertainty (Houslay and Wilson 2017). However, we used BLUPs because summarizing parental care in this way should not substantively shift nests from one parental care level to another and given the advantages of using a mixed-method model to summarize our data as described above.

### Statistical analyses

We conducted four sets of analyses, in alignment with our objectives and hypotheses (**Figure 1**), for which specific models and details are described in subsequent paragraphs. Across all analyses, we report both unadjusted as well as adjusted associations from multiple variable regression models. We selected covariates for multiple variable models based on bivariate associations and *a priori* knowledge. For all models, we z-score standardized numeric independent variables and covariates to ensure variables were modeled on the same scale. We assessed the assumptions of normality and homoscedasticity by viewing residual plots. All steps of the analyses were completed in R, version 4.2.1 (R Core Team 2023), and the ‘tidyverse’ package, version 1.3.2 was used for data cleaning and organization (Wickham et al. 2019). Models were fit using the ‘lme4’ package, version 1.1-30 (Bates et al. 2015), and when estimating marginal means, we used the ‘emmeans’ package, version 1.7.5 (Lenth 2022). For each model we estimated the marginal and conditional R-squared values using the ‘performance’ package, version 0.12.2 (Nakagawa and Schielzeth 2013; Lüdecke et al. 2021). We set α = 0.05 as the threshold for statistical significance.

#### Step 1: Characterize glucose response to an acute stressor

Our first set of analyses assessed whether an acute stressor causes an increase in blood glucose levels among nestlings at mid- (n = 104) and late development (n = 106) using linear mixed models. In these models, the explanatory variable of interest was a two-level ordinal variable indicating whether a sample was collected at baseline or following the acute stressor, and fasting blood glucose levels were included in the model as the outcome. We controlled for nest age and the sampling time as precision covariates. To account for repeated sampling of individuals and clustering by nest, we included a random intercept for nest and for nestling ID nested within nest. We first ran the model with a product term between developmental stage and time of blood collection to assess for life stage dependent effects (*i.e.,* if the statistical significance of the product term was P<0.05). Subsequently, we ran models stratified by developmental stage, as appropriate based on the P-interaction.

#### Step 2: Estimate individual repeatability in glucose measures across development

We estimated repeatability of individual glucose measures between mid- and late development for baseline and stress response levels, the latter of which was calculated as stress-induced glucose minus baseline glucose. Here, we used the ‘irrICC’ package, version 1.0 to calculate individual repeatability (Gwet 2019). We report the intraclass correlation coefficient (ICC), which is based on the variance decomposition from a one-factor ANOVA and calculated as the ratio of between subject variance divided by total variance (i.e., between subject plus within subject).

#### Step 3: Test hypotheses on the association between size/growth and glucose physiology

We tested hypotheses that nestling size and growth are associated with blood glucose levels using mixed-effects linear regression. In separate models, we estimated the effect of nestling size, measured as right-wing length, relative nestling size, and growth on fasting blood glucose, measured at baseline and following an acute stressor. In adjusted models, we controlled for nest age as a confounding variable and the time at baseline blood sampling as a precision covariate and included a random intercept for nest ID in all models. We examined cross-sectional associations at mid- and late development, as well as prospective associations of mid-development size with late development blood glucose metabolism.

#### Step 4: Assess effect modification by parental care

In our final set of models, we tested the hypothesis that the associations of nestling size and growth with blood glucose levels depend on the amount of parental care that nestlings receive. In this context, parental care is an effect modifier – i.e., a variable that introduces context dependence for the relationship of interest. Here, we focused on the relationship between morphological (nestling size and growth) and physiological variables (glucose) during late development because this was when we observed the strongest associations in prior steps and because it allowed us to assess the importance of parental care across development. To quantitatively evaluate whether the relationship between morphology and physiology differed by parental care, we included a product term between each morphology measure and parental care (low vs. high). The P-value for this product term was used to guide interpretation of results from stratified analysis of the relationship between morphological traits and physiology, within levels of parental care. We include estimates from the stratified models because they provide a direct comparison to the predictions from life history theory, in which trade-offs or differential acquisition are expected to dissipate in contexts where resources are abundant. In all models we again included a random intercept for nest ID and adjusted for the age of the nest and the time at baseline blood sampling.

#### Sensitivity analyses

We conducted several sensitivity analyses to assess robustness of estimates to differences in model parameterization and sample selection criteria. For Step 1, we included the number of nestlings as a covariate in our models assessing glucose response to handling stress in mid- and late development. We conducted another sensitivity analysis for Steps 1 and 2 by restricting the data to baseline samples collected in less than 4 minutes. For Step 3, we compared estimates from our main model to a model that also accounted for the number nestlings and a model in which baseline blood samples were collected within 4 minutes of capture.

## RESULTS

### Background characteristics

At mid-development (day 8-9), mean ± SD right-wing length was 30.71 ± 5.36 mm. During late development (day 11-13), nestlings had a right-wing length of 52.78 ± 6.44 mm. At mid-development, nestling barn swallows had a fasting baseline blood glucose level of 193.70 ± 26.31 mg/dl and stress-induced glucose level of 199.13 ± 32.36 mg/dl. In late development, nestlings had a fasting blood glucose level of 194.91 ± 30.94 mg/dl at baseline and of 230.29 ± 39.62 mg/dl following an acute stressor (**Supplemental Table 3**).

Focusing on measures of parental care, the combined feeding rates from both social nest parents were 12.6 ± 6.96, 13.0 ± 7.94 and 20.1 ± 13.0 feeding counts/hr at early, mid-, and late development, respectively (**Supplemental Table 3**). Additional background characteristics of the study population are included in **Supplemental Table 3**. Associations between covariates and size/growth measures are in **Supplemental Table 4**, and associations between covariates and blood glucose levels are in **Supplemental Table 5**.

### Step 1: Glucose increases in response to an acute stressor in late development

In nestling barn swallows, glucose response to an acute stressor depends on developmental stage (P_collection time x developmental stage_ < 0.001; marginal R^2^ =0.17, conditional R^2^ = 0.43). More specifically, at mid-development (n = 104), fasting blood glucose did not differ between baseline (reference group) and stress-induced measures (β = -0.76 [95% CI: -41.53, 40.63 mg/dl]; marginal R^2^ =0.04, conditional R^2^ = 0.66). In nestlings at late development (n = 106), fasting blood glucose increased from baseline (reference group) to stress-induced measures (β = 52.89 [95% CI: 13.94, 91.56 mg/dl]; marginal R^2^ = 0.22, conditional R^2^ = 0.74; **Figure 3**; **Supplemental Table 7**). Sensitivity analyses for number of nestlings and restricting the data to baseline samples collected in less than 4 minutes did not substantively change the direction or magnitude of the estimates of interest (**Supplemental Table 8)**.

**Figure 3.**
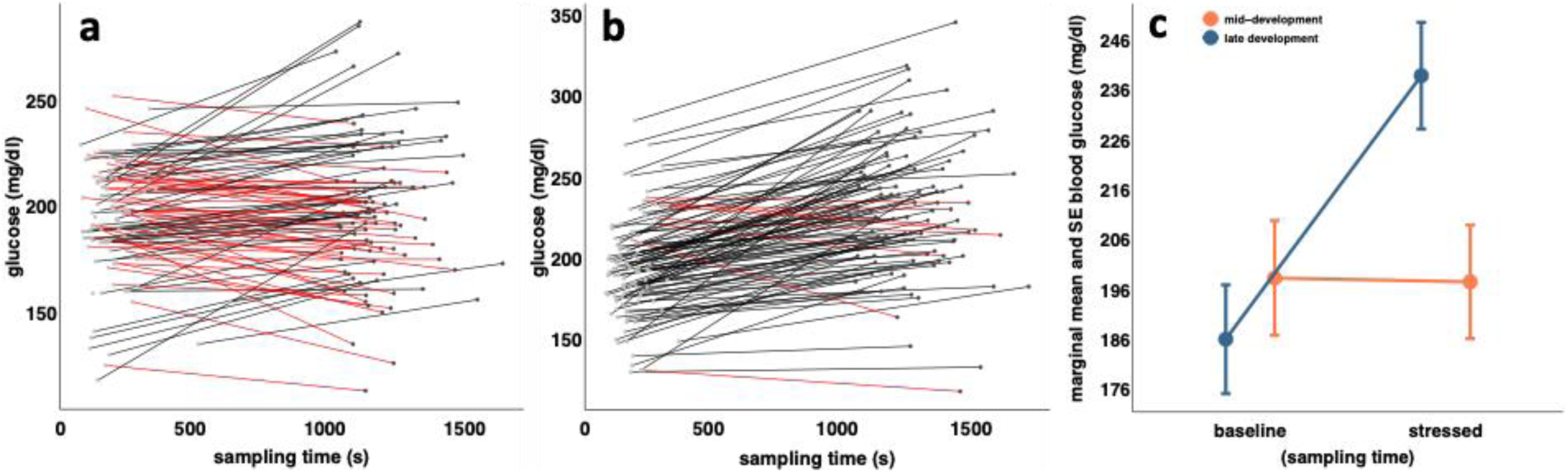
**a** Raw data for mid-development glucose measures between baseline (light gray dots) and stress response levels (dark gray dots). Each line represents the change for an individual. Red lines indicate negative change from baseline to stress response levels, and black lines correspond with a positive change. **b** Glucose measures for the same individuals at late development. **c** A comparison of the glucose response measured as the difference between baseline and stress response levels assessed in nestlings during both mid-development (orange) and late development (blue). The dot and whiskers correspond to the estimated marginal means of each group and the corresponding standard errors.

### Step 2: Baseline but not stress response blood glucose exhibit low to moderate repeatability over development

We observed mixed results for intraindividual repeatability in blood glucose levels. The intraclass correlation coefficient (ICC) for baseline glucose levels across mid- and late development was 0.29 (95% CI: 0.09, 0.45) while the ICC for the stress response glucose levels across mid- and late development was 0.00 (95% CI: 0.00, 0.02; **Supplemental Table 9**). When we restricted the data to baseline samples collected in less than 4 minutes, we observed no substantive changes in the direction, magnitude, or precision of the estimates of interest (**Supplemental Table 10**).

### Step 3: Nestling size and growth are positively associated with glucose measures

At mid-development, longer right-wing length was associated with higher baseline glucose levels (6.00 [95% CI: 0.29, 11.69] mg/dl per 1 SD wing length; marginal R^2^ = 0.05, conditional R^2^ = 0.33). The effect of relative nestling size was in the same direction, with larger nestlings having higher baseline glucose levels than smaller nestlings (7.49 [95% CI: -1.55, 16.65] mg/dl; marginal R^2^ = 0.03, conditional R^2^ = 0.31 **Table 1**), although this effect was not significant. There was no effect of nestling size on stress response glucose levels at mid-development (**Table 1**).

**Table 1.** Associations between size and growth and fasting blood glucose levels assessed in nestling barn swallows during mid- and late-development.

| Models | Mid-development (~8 days post hatch) |  |  |  | Late-development (~12 days post-hatch) |  |  |  |
| --- | --- | --- | --- | --- | --- | --- | --- | --- |
|  | Baseline glucose (mg/dl) |  | Glucose response to handling stress (mg/dl) <sup>a</sup> |  | Baseline glucose (mg/dl) |  | Glucose response to handling stress (mg/dl) <sup>a</sup> |  |
| | N | $\beta$ (95% CI) <sup>b</sup> | N | $\beta$ (95% CI) <sup>b</sup> | N | $\beta$ (95% CI) <sup>b</sup> | N | $\beta$ (95% CI) <sup>b</sup> |
| **Mid-development right-wing length (z-score standardized)** |  |  |  |  |  |  |  |  |
| unadjusted | 104 | 5.82 (0.67, 10.97) | 100 | 2.12 (-2.88, 7.02) | 103 | 6.46 (0.65, 12.18) | 103 | 6.78 (1.11, 12.46) |
| adjusted <sup>c</sup> | 104 | 6.00 (0.29, 11.69) | 100 | 0.10 (-5.44, 5.65) | 103 | 8.13 (2.44, 13.72) | 103 | 4.48 (-1.44, 10.17) |
| **Mid-development relative size <sup>d</sup>** |  |  |  |  |  |  |  |  |
| unadjusted | 104 | 7.62 (-1.43, 16.75) | 100 | -3.31 (-13.17, 6.67) | 103 | 7.26 (-2.45, 16.94) | 103 | 0.45 (-11.26, 11.69) |
| adjusted <sup>c</sup> | 104 | 7.49 (-1.55, 16.65) | 100 | -3.69 (-13.37, 6.00) | 103 | 8.62 (-0.69, 17.91) | 103 | -1.36 (-12.85, 9.65) |
| **Late-development right-wing length (z-score standardized)** |  |  |  |  |  |  |  |  |
| unadjusted | - | - | - | - | 106 | 9.33 (4.32, 15.55) | 106 | 8.49 (2.78, 14.20) |
| adjusted <sup>c</sup> | - | - | - | - | 106 | 11.16 (5.47, 17.00) | 106 | 6.55 (0.25, 12.63) |
| **Late-development relative size <sup>d</sup>** |  |  |  |  |  |  |  |  |
| unadjusted | - | - | - | - | 106 | 3.14 (-6.18, 12.46) | 106 | 7.14 (-3.61, 17.74) |
| adjusted <sup>c</sup> | - | - | - | - | 106 | 4.32 (-4.61, 13.31) | 106 | 6.40 (-4.52, 16.87) |
| **Growth based on right wing-length difference (z-score standardized)** |  |  |  |  |  |  |  |  |
| unadjusted | - | - | - | - | 103 | 8.37 (1.91, 14.88) | 103 | 4.45 (-1.87, 10.81) |
| adjusted <sup>c</sup> | - | - | - | - | 103 | 7.32 (0.66, 14.24) | 103 | 4.01 (-2.45, 10.24) |

<sup>a</sup> Glucose response to handling stress is calculated as the difference between the stressed state concentration minus the baseline concentration of fasting blood glucose levels.<sup>b</sup> Estimated $\beta$ (95% CI) from linear mixed models in which nest ID was included as a random intercept.<sup>c</sup> Adjusted models include the age of the nest, as determined from the day that the first nestling hatched, and the time (s) elapsed between nestling removal from the nest and the blood draw.<sup>d</sup> Relative size is calculated based on an individual's right wing length below (small) or above (large) the average right wing length within a nest. Small is set as the reference category.Bold estimates are significant at $P < 0.05$ .

During late development, we also observed that larger nestlings, as determined by right-wing length, had higher baseline glucose (11.16. [95% CI: 5.47, 17.00] mg/dl per 1 SD wing length; marginal R^2^ = 0.15, conditional R^2^ = 0.51). Greater growth between mid- and late development was also associated with higher baseline blood glucose (7.32 [95% CI: 0.66, 14.24] mg/dl per 1 SD difference in wing length; marginal R^2^ = 0.11, conditional R^2^ = 0.46) during late development. The largest nestlings based on relative size again had higher baseline blood glucose (4.32 [95% CI: -4.61, 13.31] mg/dl; marginal R^2^ = 0.06, conditional R^2^ = 0.48), though this effect was not significant. In late development, larger right-wing length corresponded with an augmented glucose stress response (an increase of 6.55 [95% CI: 0.25, 12.63] mg/dl per 1 SD in wing length; marginal R^2^ = 0.13, conditional R^2^ = 0.24). Larger relative nestling size and greater growth were both positively associated with late development stress response glucose levels, but these estimates were not significant (**Table 1**).

In sensitivity analyses, adjustment for the number of nestlings and restricting the data to baseline samples collected in less than 4 minutes did not substantively change the direction, magnitude, or precision of the estimates of interest (**Supplemental Table 11**).

### Step 4: Evidence of differential allocation in late development is limited to nests that received low parental care

The interaction between right-wing growth and parental care with glucose response approached significance (P = 0.08; marginal R^2^ =0.11, conditional R^2^ = 0.40), but in none of the other three interaction models were the product terms involving measures of size/growth and parental care significant (**Supplemental Table 12)**. However in stratified analysis, longer right-wing length corresponded with higher baseline glucose among nestlings that received low parental care (15.38 [95% CI: 6.25, 24.90] mg/dl per 1 SD wing length; marginal R^2^ =0.29, conditional R^2^ = 0.39) but we observed no association in high parental care nests (4.70 [95% CI: -3.17, 13.27] mg/dl per 1 SD wing length; marginal R^2^ =0.12, conditional R^2^ = 0.60; **Figure 4a**). Similarly, greater growth was associated with higher baseline glucose among nestlings that received low parental care, although this association was not significant (8.43 [95% CI: -0.18, 16.96] mg/dl per 1 SD growth; marginal R^2^ =0.19, conditional R^2^ = 0.35). Growth was not associated with baseline glucose among nestlings from high parental care nests (-2.08 [95% CI: -14.25, 12.47] mg/dl per 1 SD growth; marginal R^2^ =0.09, conditional R^2^ = 0.64; **Figure 4b**). Both longer right-wing length and greater growth were associated with higher stress response glucose levels among nestlings that received low parental care, 12.53 (95% CI: 2.51, 22.31; marginal R^2^ =0.13, conditional R^2^ = 0.15; **Figure 4c**) mg/dl and 12.10 (95% CI: 2.53, 20.52; marginal R^2^ =0.16, conditional R^2^ = 0.26; **Figure 4d**) mg/dl, respectively. However, the associations between right-wing length (4.17 [95% CI: -5.05, 14.45]; marginal R^2^ =0.32, conditional R^2^ = 0.57; **Figure 4c**) and growth (-2.81 [95% CI: -15.65, 10.10]; marginal R^2^ =0.25, conditional R^2^ = 0.58; **Figure 4d**) with stress response glucose levels were attenuated towards the null when restricted to nestlings from high parental care nests.

**Figure 4.**
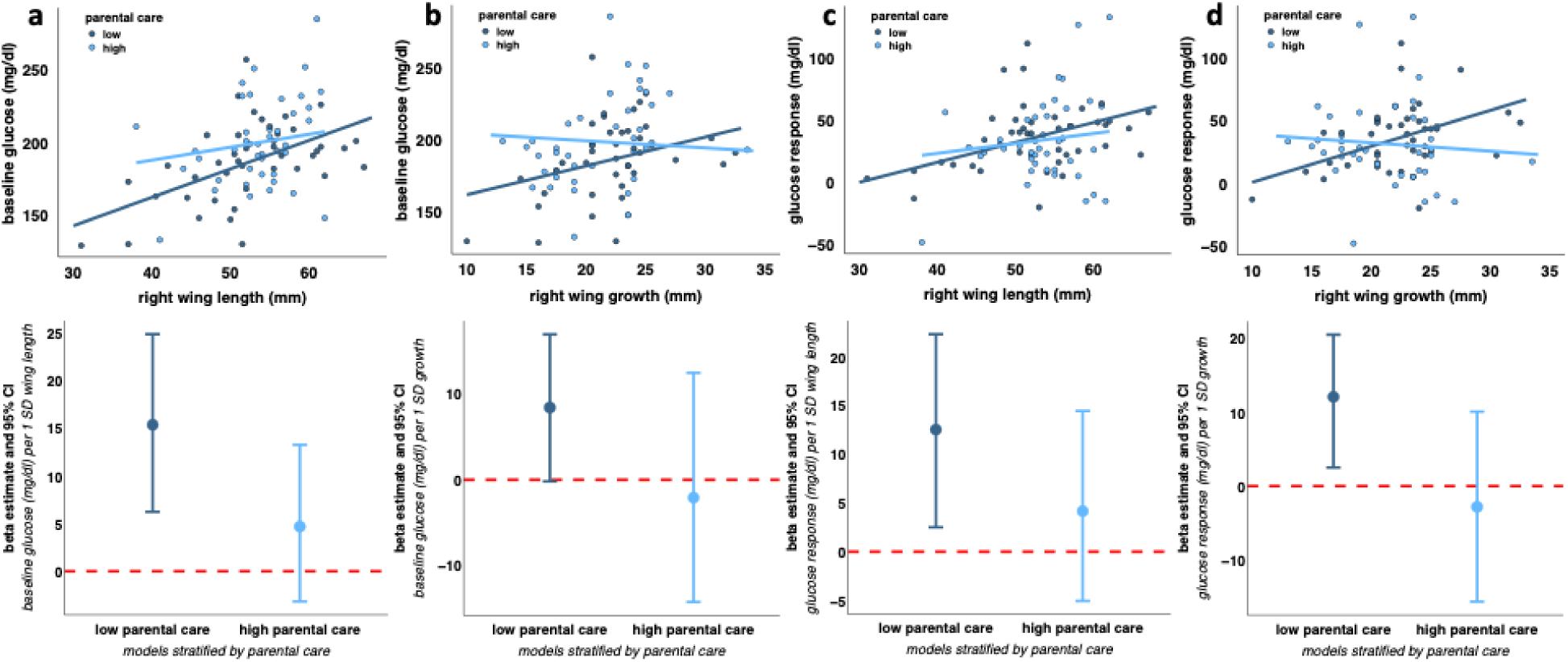
In the top row are the raw data for morphological and physiological measures in nestling barn swallows that received low (dark blue) or high (light blue) parental care. The corresponding color matched lines are the predicted values from separate models stratified by level of parental care. In the bottom row are beta estimates and 95% CIs for the relationship between size/growth and blood glucose levels from models stratified by level of parental care. **a** This column corresponds to the association between right-wing length and baseline glucose. **b** This column corresponds to the association between growth and baseline glucose. **c** This column corresponds to the association between right-wing length stress response glucose. **d** This column corresponds to the association between growth and stress response glucose. Right-wing length and blood glucose levels were assessed in late development.

## DISCUSSION

In this study of 113 barn swallow nestlings, we characterized baseline and stress response fasting blood glucose levels as indicators of physiological condition across development. We also investigated associations of nestling size and growth with these endpoints to understand life history strategies surrounding early-life growth and physiology. We observed an increase fasting blood glucose in response to an acute stressor in late development but not mid-development. This life-stage dependent association is reflected by the low to moderate intraindividual repeatability in baseline and a lack of repeatability in stress response glucose levels between mid- and late development in this sample. We also found that larger nestlings had higher baseline glucose levels than their smaller counterparts, and a more pronounced glucose response to an acute stressor in late development. Finally, we found evidence of effect modification by parental care such that birds from nests that received the lowest parental care showed the strongest associations of size and growth with baseline glucose and change in glucose levels before vs. after an acute stressor. This pattern suggests that parental failure to adequately provision food for nestlings, can exacerbate unequal resource acquisition such that smaller nestlings get fewer resources and larger nestlings get more.

### Glucose increases in response to an acute stressor in late development

Our finding that barn swallows exhibit an increase in blood glucose in response to an acute stressor during late development conforms with the mechanistic explanation that HPA axis activation mobilizes glucose into the bloodstream (Sapolsky et al. 2000). Vertebrates are adapted to deal with acute stressors by allocating energy to overcome and recover from challenges (McEwen and Wingfield 2003; Romero et al. 2009; Romero and Beattie 2022). This process is facilitated on the order of seconds to minutes by secretion of catecholamines, and on the order of minutes to hours by glucocorticoids, including corticosterone, both of which result in higher blood glucose concentrations (Sapolsky et al. 2000; Romero and Beattie 2022). Consistent with our results are findings from studies in birds and reptiles confirming a general pattern in which blood glucose levels increase following exposure to acute stressors, such as handling and restraint during capture (Remage-Healey and Romero 2000, 2001; Gangloff et al. 2017; Montoya et al. 2020; Neuman-Lee et al. 2020; Vágási et al. 2020; Taff et al. 2022).

Acute stress exposure does not always result in an increase in blood glucose levels. For example, male Rufous-winged Sparrows (*Peucaea carpalis*) exhibit a stress-induced increase in blood glucose levels prior to their breeding season, no response in glucose during their breeding season and a negative glucose response to stress post-breeding (Deviche et al. 2016). Moreover, some studies in birds (Taff et al. 2022) and reptiles (Gangloff et al. 2017; Neuman-Lee et al. 2020) that measured both circulating glucocorticoids and glucose noted a stress-induced increase in glucocorticoids does not correlate with an increase in glucose. A lack of correlation between circulating glucocorticoids and glucose in response to acute stressors underscores the complexity in the regulation and timing of physiological pathways that control downstream metabolites that can be measured in blood. The influence of both external and internal factors on the function of the physiological mechanisms involved in stress response highlight that additional work is needed to both understand the proximate mechanisms that allow organisms to cope with acute stressors and to ultimately link an individual’s physiological stress response with their fitness (Romero and Gormally 2019).

Of note, we did not observe a stress-induced glucose response during mid-development, suggesting that at this younger age, barn swallows may be physiologically limited in their capacity to respond to an acute stressor due to small size and/or young age. Indeed, studies in several other avian species observed that post-hatch HPA activity, as measured by baseline or stress-induced corticosterone levels, was larger in older nestlings (Schwabl 1999; Sims and Holberton 2000; Love et al. 2003). In a recent study of a closely related species of tree swallows (*Tachycineta bicolor)*, Taff *et al*. reported a similar pattern of no significant change in blood glucose before vs. after a stressor among nestlings but not adults (Taff et al. 2022). Similarly, northern mockingbirds (*Mimus polyglottos*) show an age dependent capacity to increase circulating glucocorticoids in response to handling stress that suggests limited functional capacity of the HPA axis early in life (Sims and Holberton 2000). We conclude that a physiological response to an acute stressor appears limited in very young or small swallows, and that the ability to increase blood glucose and mobilize energy is a developmentally constrained adaptation that is likely important as nestlings near fledging.

### Baseline but not stress response blood glucose has moderate repeatability over development

We observed moderate intraindividual repeatability in baseline blood glucose levels, with comparable ICC to that reported in zebra finches (*Taeniopygia guttata;* ICC ∼0.3) (Montoya et al. 2018). On the other hand, ICC for the stress response glucose levels across mid- and late development was null (ICC ∼0.0) reflecting substantial variation in capacity to mount a physiological response to stressors across development. This finding is likely related to results from Step 1 in which we observed a glucose response to stress during late but not mid-development. Despite a general pattern of moderate repeatability in blood glucose measurements among various bird species, all reported ICC values fall below the level of 0.50, indicating relatively high within-individual variation in glucose measures (Koo and Li 2016; Taff et al. 2022). Such variation could reflect individual exposures to heterogenous environments, especially if glucose metabolism is a plastic process.

### Nestling size and growth are positively associated with glucose measures

In support of the differential acquisition hypothesis (**Figure 1**), we found that larger nestlings and those that grew faster had higher baseline glucose levels at mid- and late development than smaller nestlings. Higher baseline glucose may fuel activity of larger nestlings as they prepare to fledge given that glucose is an important source of energy (Braun and Sweazea 2008). Higher baseline glucose may also provide developing nestlings with essential energy for thermoregulation (Montoya et al. 2018; Sweazea et al. 2020). Regardless, we acknowledge studies in blue tits (*Cyanistes caeruleus*), great tits (*Parus major*) and zebra finches have reported inverse (Kaliński et al. 2014, 2015) or null (Montoya et al. 2018; Kaliński et al. 2019) associations of mass and size with baseline blood glucose levels. A study comprising multiple species of birds found both negative and positive associations between mass and baseline blood glucose in adult, grey-breasted white-eyes (*Zosterops lateralis*) and welcome swallows (*Hirundo neoxena*), respectively (Lill 2011). The same study also found that in welcome swallows and spotted doves (*Spilopelia chinensis*), baseline blood glucose was higher in later development when nestlings are older and heavier (Lill 2011). An increase in circulating glucose concentrations with increasing age across early development has also been observed in poultry (Lu et al. 2007; Richards et al. 2010). Positive, negative, and null associations between structural and physiological traits may reflect species-specific life history strategies involving difference in allocation versus acquisition of resources. Associations between size and baseline blood glucose are expected to differ across ontogeny (Lu et al. 2007; Richards et al. 2010; Lill 2011) if there are developmental constraints on physiology, or by environment (Montoya et al. 2018), especially if the trait exhibits plasticity.

We also found that larger nestlings exhibited a greater glucose response to an acute stressor in late development. Capacity to mount a robust glucose response may improve survival at fledging by mobilizing energy that allows nestlings to escape and recover from interactions with predators or to cope with environmental challenges encountered outside of the nest. A prediction following this functional explanation is that size as well as proxies for size (e.g., hatch order) correspond to differences in stress response corticosterone levels. Love *et al*. report that first hatched and larger nestling American kestrels (*Falco sparverius*) show a stronger stress-induced corticosterone response than later hatched siblings (Love et al. 2003). Their result parallels the greater increase in stress-induced glucose that we observe in larger nestling barn swallows during late development. Such positive trait associations lend support to the hypothesis that larger nestlings are capable of a more robust physiological response, potentially because they acquire more resources and are in better condition. This stands in contrast to negative associations between structural and physiological traits which better aligns with the differential allocation hypothesis involving a trade-off between competing traits (Van Noordwijk and De Jong 1986; Stearns 1989).

### Evidence of differential allocation is limited to birds from nests that receive low parental care

Finally, following the prediction that constraints are revealed when resources are limited, we found that while larger and faster-growing nestlings had higher baseline glucose and a greater magnitude of response to an acute stressor, these associations were strongest among nestlings in nests that received the least amount of parental care. These results suggest that inadequate parental care may exacerbate the limited physiological capacity of the smallest nestlings. Such effect modification by parental care on the relationship between offspring size and development has been reported in other study systems as well. For example, a study in burying beetles (*Nicrophorus vespilloides*) found that larger egg size corresponded with larger post hatch body mass, but only in the absence of parental care (Monteith et al. 2012). Accordingly, in this study, parental care buffered any disadvantage associated with small egg size, a pattern that parallels our results. We posit that when inadequate parental care limits resources available within a nest, there will be greater variation in the distribution of resources among nestlings. Specifically, in the absence of pronounced trade-offs, theory predicts a positive association between structural and physiological traits (Van Noordwijk and De Jong 1986; Stearns 1989), with larger individuals and those in better condition obtaining more resources than their smaller, less physically robust counterparts. Future studies with more detailed measures of resource acquisition, particularly in nests with low parental care, could improve future hypothesis testing regarding how differential allocation versus acquisition of resources affects life history traits, especially if individual nestling behaviors influence the distribution of resources (Laskowski et al. 2021).

### Strengths and limitations

This study has several strengths. First, in a relatively large sample size of nestling barn swallows (n =113), we measured fasting blood glucose levels during the early morning. Through this sampling design we reduced potential confounding since circulating glucose levels are also regulated by insulin and other peptide hormones that are influenced by circadian rhythms and food intake (Remage-Healey and Romero 2000, 2001; Kuo et al. 2015). Second, we measured blood glucose levels in the first part of the breeding season in first broods and in a single year, thereby limiting extraneous variation in glucose due to seasonal and yearly fluctuations in the environment (Remage-Healey and Romero 2000; Kaliński et al. 2014, 2015; Montoya et al. 2018). Third, we collected data longitudinally, which allowed us to measure baseline and glucose response to an acute stressor in the same individuals during different phases of development. This strengthens our evidence that young birds are unable to increase circulating glucose when exposed to perturbations because they are developmentally constrained. The longitudinal data also allowed us to assess individual level differences in size and physiology. Finally, we were able to combine detailed parental care data collected over the course of nestling development with nestling morphology and physiology data, allowing us to test the hypothesis that early-life environmental factors limit resource access affecting the relationship between structural and physiological trait development.

Our study was not without limitations. First, we lacked data on conventional stress hormone biomarkers (*e.g.*, corticosterone), which limited our ability to make inference on the physiological pathways involved in glucose regulation at baseline and as part of the HPA stress response mechanism (Vágási et al. 2020). We were additionally limited by the ability to collect enough blood volume for multiple biomarkers, and in this study, we focused on dividing small blood samples into amounts for glucose measures and storage in lysis buffer for later DNA extraction. Second, since we did not mark individuals upon hatching, age and size cannot be differentiated in our study. While we discuss in detail ontogenetic explanations for our results, being able to assess whether it was age or size *per se* that affects glucose levels would be informative. Third, while we interpreted higher baseline fasting glucose and a larger change in glucose before vs. after exposure to a stressor as favorable physiological outcomes, this may not always be the case given that persistently and excessively high glucose levels can lead to oxidative stress and cell damage. However, without established thresholds for what constitutes “high” blood glucose levels in avian species, and in light of evidence that blood glucose levels within the range of those observed in the present analysis occur without causing oxidative or vascular damage, we do not anticipate this being a major concern for this study (Braun and Sweazea 2008; Smith et al. 2011; Scanes and Braun 2013; Sweazea 2022).

To address the above limitations, we suggest future studies 1) keep individual level records on both age and size, 2) consider an experimental manipulation of hatch asynchrony (see a study in pied flycatchers (*Ficedula hypoleuca*) (Kärkkäinen et al. 2021) to improve causal inference on the effect of structural traits on physiological development, and 3) implement a nest-cross fostering treatment to limit potential reverse causation in which nestling traits cause differences in parental behavior. Finally, while we observed statistically significant associations in this study, future studies with even larger sample sizes will allow for detection of smaller yet biologically relevant effect sizes between early growth and blood glucose levels that may track across the life span.

### Conclusion

In this study of wild barn swallow nestlings, we found that larger nestlings and those that grew faster exhibited the most favorable glucose metabolism outcomes later in development. Further, these associations were modified by parental care such that inadequate care exacerbated the detrimental effect of being small and growing slowly.

Viewed through a life history lens, these findings are indicative of differences in resource acquisition. Although it has long been theorized that differences in resource allocation versus differences in resource acquisition affect the covariation of life history traits (Van Noordwijk and De Jong 1986; Stearns 1992), recognition that within species and among individual trade-offs can be masked by variation in resource acquisition has garnered recent interest (Laskowski et al. 2021). This study provides empirical evidence supporting the hypothesis that larger differences in resource acquisition relative to difference in resource allocation can result in positive trait covariation and veil trade-offs if they exist. Our findings also suggest that parental behaviors can influence resource acquisition, making it an important contextual variable to consider when testing life history hypotheses (Laskowski et al. 2021). A promising direction for future research is longitudinal studies that can elucidate how patterns of growth and development of physiology early in life under different levels of parental care affect fitness.

## Data accessibility

All data and analysis code used in this paper are available at this public GitHub repository https://github.com/laubach/bs_parent_care_modifies_size_phys.git.

## Author contributions

Z.M.L. and R.J.S. conceptualized research. Z.M.L., S.A.M., and A.P. collected data; Z.M.L. conducted analyses; Z.M.L. wrote original draft. All authors contributed critically to the drafts and gave final approval for publication.

## Acknowledgements

We thank Kim Hoke and Wei Perng for thoughtful feedback on early drafts of this manuscript. We also thank Marina Ayala, Heather Kenny-Duddela, Avani Fachon, Grant Gonzalez, and Molly McDermott for assistance and/or advice regarding field work. We thank the private landowners who give us access to their property where barn swallows nest. Finally, we thank members of the Safran and Hoke labs for their constructive feedback on this project.

