## Supplemental Material for "Parental care modifies the role of early-life size and growth in shaping future physiology"

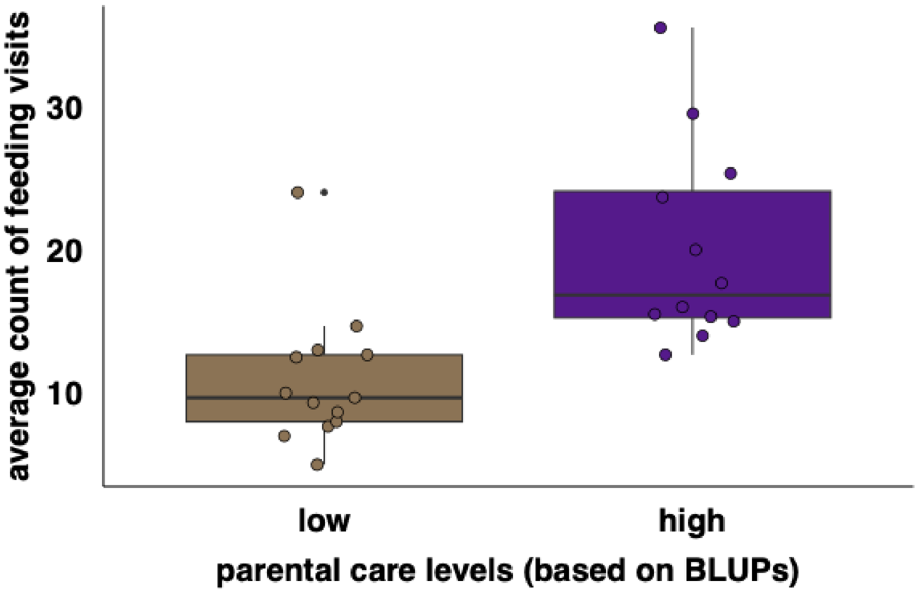


**Figure S1.** Histograms of counts feeding visits for each nest averaged across early, mid-, and late development by level of parental care. Parental care levels are based on Best Linear Unbiased Predictors of feeding counts for each nest, split below, low parental care, and above, high parental care, the median.

**Supplemental Table 1**: Ethogram of parental care behaviors.

| Behavior location | Behavior name | Behavior type | Description | Developmental context notes |
|---|---|---|---|---|
| At nest | Feeding | Event | The parent delivers food as indicated by putting its beak into the mouth of the nestling. Prey often cannot be seen in the parent's bill. Feeding usually occurs immediately after arrival at the nest. | Day 3: nestlings are small, so they usually cannot be seen by the observer. Feeding visits are identified by the parent leaning into the nest immediately after arrival from foraging.<br>Day 8: the parent can often be seen inserting its bill into the gap of nestlings. The parent usually still leans into the nest slightly when feeding.<br>Day 12: nestlings can be seen and feeding occurs without leaning into the nest. |
| | Brooding | State | The parent settles itself all the way into the nest on top of the nestlings. The parent may turn around in the nest while brooding. Brooding behavior looks similar across nestling ages but is far more common with younger nestlings (Day 3). | NA |
| | Sanitizing | State | The parent removes nestling fecal sacs or rearranges nestlings and feathers in the nest. The parent may carry fecal sacs out of the nest or consume them. When rearranging feathers or nestlings, the parent may perch on the edge of the nest and lean in deeply, tail twitching (see-saw motion; head and tail pivot up and down). The parent may also sanitize while brooding, as indicated by their head in the nest and tail twitching. | Day 3: The parents often sanitize while settled in the nest cup. The parent will shift around and the tail will twitch. The head will duck down into the nest.<br>Days 8 and 12: Sanitizing more common and occurs while parent is perched on the nest edge. Parents may be seen carrying fecal sacks away. |
| | Preening | State | The parent uses the beak to clean, stroke, or re-position its feather while perched on the nest rim or settled in the nest brooding. | NA |
| | Perching | State | The parent stands with feet on the side of the nest, not engaged in any other behaviors. The parent may look around the barn or it may peer inside the nest. | NA |
| Away from nest | Near nest | State | The parent is visible at their typical perch location(s). Importantly, here it is reasonable to assume that the bird can see the nest (i.e., they are able to be vigilant of any nest threats). | NA |
| | Far from nest | State | The parent flies away from the nest and is either observed leaving the barn or headed towards a barn opening but that might be out of sight for the observer. | NA |
| | Unkown | State | It is not known if the parent can reasonably see the nest or is in the barn at all. | NA |

**Supplemental Table 2.** AIC comparison of models used to generate the parental care feeding BLUPs for different distrubutions.

| Model distribution | dAIC | Degrees of freedom |
|---|---|---|
| negative binomial | 0 | 7 |
| Poisson | 18.2 | 6 |
| Gaussian | 21.8 | 7 |

Covariates in each all mixed models included: the number of days since the first nestling hatched, the number of nestlings in a nest, the median temperature near the nest during the observation trial, and the duration of time between extracting nestlings from the nest and the start of the focal data collection. The outcome was repeated measures of counts of feeding visits from both parents. Each model also included a random intercept for nest ID and an offset for the total time a nest was observed.

The marginal $R^2$ = 0.33 and the conditional $R^2$ = 0.44 from final negative binomial model.

Best Linear Unbiased Predictors (BLUPs)

**Supplemental Table 3.** Background characteristics of nestling swallows summarized over the course of development.

| | Early development (~3 days post hatch) | | Mid-development (~8 days post hatch) | | Late development (~12 days post hatch) | |
|---|---|---|---|---|---|---|
| | *N* | mean ± SD | *N* | mean ± SD | *N* | mean ± SD |
| **Blood glucose levels (mg/dl)** | | | | | | |
| Baseline | - | - | 104 | 193.70 ± 26.31 | 106 | 194.91 ± 30.94 |
| Stress induced | - | - | 100 | 199.13 ± 32.36 | 106 | 230.29 ± 39.62 |
| **Size and mass** | | | | | | |
| Right wing length (mm) | 113 | 9.40 ± 1.94 | 108 | 30.71 ± 5.36 | 106 | 52.78 ± 6.44 |
| Growth (late - mid right wing length, mm) | - | - | - | - | 103 | 21.77 ± 4.25 |
| **Parental care behavior** [a] | | | | | | |
| Total feeding rate (counts/per hr) | 24 | 12.6 ± 6.96 | 22 | 13.0 ± 7.94 | 23 | 20.1 ± 13.0 |

[a] Parental care behaviors are measured at the level of the nest and include the totals for both social parents (maternal and paternal care). Total nest feeding is reported as a rate (counts/observation time).

**Supplemental Table 4.** Background characteristics of nestling swallows summarized by level of parental care.

| | Mid-development (~8 days post hatch) | | | | Late development (~12 days post-hatch) | | | |
|---|---|---|---|---|---|---|---|---|
| | Low parental care [a] | | High parental care [a] | | Low parental care [a] | | High parental care [a] | |
| | *N* | mean ± SD | *N* | mean ± SD | *N* | mean ± SD | *N* | mean ± SD |
| **Blood glucose levels (mg/dl)** | | | | | | | | |
| Baseline | 46 | 191.70 ± 26.04 | 36 | 195.53 ± 21.46 | 42 | 186.81 ± 27.61 | 42 | 200.10 ± 30.19 |
| Stress induced | 43 | 199.05 ± 36.49 | 36 | 198.58 ± 18.68 | 42 | 222.55 ± 36.95 | 42 | 233.86 ± 38.69 |
| **Size and mass** | | | | | | | | |
| Right wing length (mm) | 46 | 29.83 ± 5.36 | 39 | 32.27 ± 5.17 | 42 | 52.33 ± 7.85 | 42 | 53.57 ± 5.26 |
| Growth (late - mid right wing length, mm) | - | - | - | - | 42 | 22.21 ± 4.23 | 39 | 21.13 ± 4.21 |

[a] Parental care behaviors are measured at the level of the nest and include the totals for both social parents (maternal and paternal care) summarzied across development using mixed models to extract nest-level Best Linear Unbiased Predictos (BLUPs) and then categorized dichotomously.

**Supplemental Table 5.** Associations between barn swallow nestling covariates and measures of size and growth assessed at mid- and late development.

| | **Mid-development (~8 days post hatch)** | | **Late development (~12 days post-hatch)** | | | |
|---|---|---|---|---|---|---|
| | **Right-wing length** | | **Right-wing length** | | **Growth (difference in right-wing length) [a]** | |
| | ***N*** | ***β (95% CI)* [b]** | ***N*** | ***β (95% CI)* [b]** | ***N*** | ***β (95% CI)* [b]** |
| **Potential confounders** | | | | | | |
| Number of nestlings | 108 | -1.16 (-2.81, 0.50) | 106 | **-2.30 (-4.32, -0.28)** | 103 | -1.20 (-2.68, 0.27) |
| Mites (y/n) [c] | 108 | -1.14 (-4.30, 2.02) | 106 | **4.01 (0.68, 7.34)** | 103 | 1.48 (-0.70, 3.67) |
| Nest age (days since hatch) [d] | 108 | **3.51 (0.63, 6.38)** | 106 | **6.28 (3.19, 9.36)** | 103 | **3.34 (0.75, 5.92)** |

[a] Growth is calculated as the difference in right wing length between late development minus mid-development measures.

[b] Estimated β (95% CI) from linear mixed models in which nest ID was included as a random intercept.

[c] Mites are assessed on each nestling and scored as a binary variable as present (yes) or absent (no) and modeled with 'no mites' as the reference group.

[d] Nest age is based on the date when the first egg hatched.

Bold estimates are significant at $P < 0.05$.

**Supplemental Table 6.** Associations between barn swallow nestling covariates and blood glucose levels assessed at mid and late development.

| | **Mid-development (~8 days post hatch)** | | | | **Late development (~12 days post-hatch)** | | | |
|---|---|---|---|---|---|---|---|---|
| | **Glucose response to handling stress (mg/dl) [a]** | | **Baseline glucose (mg/dl)** | | **Glucose response to handling stress (mg/dl) [a]** | | **Baseline glucose (mg/dl)** | |
| | ***N*** | ***β (95% CI)* [b]** | ***N*** | ***β (95% CI)* [b]** | ***N*** | ***β (95% CI)* [b]** | ***N*** | ***β (95% CI)* [b]** |
| **Potential confounders** | | | | | | | | |
| Number of nestlings | 100 | 0.46 (-6.11, 7.03) | 104 | 3.74 (-4.38, 11.87) | 106 | -5.31 (-13.30, 2.69) | 106 | 1.29 (-8.72, 11.30) |
| Mites (y/n) [c] | 100 | 1.43 (-13.40, 16.27) | 104 | -0.49 (-16.52, 15.54) | 106 | 8.97 (-6.24, 24.18) | 106 | 6.57 (-9.64, 22.77) |
| Nest age (days since hatch) [d] | 100 | **11.93 (1.54, 22.32)** | 104 | 5.47 (-9.62, 20.55) | 106 | **13.68 (0.21, 27.14)** | 106 | 4.91 (-12.77, 22.59) |
| **Precision variables** | | | | | | | | |
| Time until baseline blood draw | 100 | -0.01 (-0.07, 0.04) | 104 | -0.02 (-0.08, 0.05) | 106 | **-0.12 (-0.21, -0.02)** | 106 | **0.12 (0.03, 0.20)** |
| Trial temperature (℃) [e] | 93 | 1.46 (-1.00, 3.93) | 96 | -0.38 (-3.84, 3.09) | 106 | 0.02 (-3.46, 3.51) | 106 | 0.84 (-3.44, 5.12) |

[a] Glucose response to handling stress is calculated as the difference between the stressed state concentration minus the baseline concentration of fasting blood glucose levels.

[b] Estimated β (95% CI) from linear mixed models in which nest ID was included as a random intercept.

[c] Mites are assessed on each nestling and scored as a binary variable as present (yes) or absent (no) and modeled with 'no mites' as the reference group.

[d] Nest age is based on the date when the first egg hatched.

[e] Trial temperature was measured at approximately sunrise on the day nestlings were handled.

Bold estimates are significant at $P < 0.05$.

**Supplemental Table 7.** Fasting glucose response to handling stress at mid- and late development in nestling barn swallows.

| | **Mid-development glucose response [a]** | | **Late development glucose response [a]** | |
|---|---|---|---|---|
| | ***N*** | ***β (95% CI)* [b]** | ***N*** | ***β (95% CI)* [b]** |
| **Model** | | | | |
| Unadjusted model | 104 | **5.20 (0.39, 10.00)** | 106 | **35.39 (29.76, 41.01)** |
| Adjusted model [c] | 104 | -0.76 (-41.53, 40.63) | 106 | **52.89 (13.94, 91.56)** |

[a] Glucose response to handling stress is estimated from models in which baseline glucose is set as the reference category for comparing the difference between stress-induced and baseline levels.

[b] Estimated β (95% CI) from linear mixed models in which nest ID was included as a random intercept and nestling ID is nested within nest.

[c] Adjusted models include the age of the nest, as determined from the day that the first nestling hatched, and the time (s) elapsed between nestling removal from the nest and the blood draw.

Bold estimates are significant at $P < 0.05$.

**Supplemental Table 8.** Sensitivity analyses for models of fasting glucose response to handling stress at mid- and late development in nestling barn swallows.

| | Mid-development glucose response [a] | | Late development glucose response [a] | |
|---|---|---|---|---|
| | *N* | *β (95% CI)* [b] | *N* | *β (95% CI)* [b] |
| **Model** | | | | |
| Adujsted for number nestlings [c] | 104 | 3.54 (-37.97, 47.58) | 106 | **53.73 (14.27, 92.60)** |
| Glucose measured in < 4min [d] | 85 | -5.19 (-68.82, 59.31) | 96 | **59.16 (15.87, 102.01)** |

[a] Glucose response to handling stress is estimated from models in which baseline glucose is set as the reference category for comparing the difference between stress-induced and baseline levels.

[b] Estimated β (95% CI) from linear mixed models in which nest ID was included as a random intercept and nestling ID is nested within nest. Models include the age of the nest, as determined from the day that the first nestling hatched, and the time (s) elapsed between nestling removal from the nest and the blood draw.

[c] Sensitivity analysis in which model is adjusted for the number of nestlings in a nest.

[d] Sensitivity analysis in which model is based on restricted data set that includes only baseline glucose measures collected in under 4 minutes.

Bold estimates are significant at $p < 0.05$.

**Supplemental Table 9.** Intraindividual repeatability (intraclass correlation coefficient) in baseline and stress response glucose levels assessed between mid- and late development.

| | Baseline glucose | | Glucose response to handling stress (mg/dl) [a] | |
|---|---|---|---|---|
| | *N* | *ICC (95% CI)* | *N* | *ICC (95% CI)* |
| **Mid- to late development** | 111 | 0.29 (0.09, 0.45) | 111 | 0.00 (0.00, 0.02) |

[a] Glucose response to handling stress is calculated as the difference between the stress-induced level minus the baseline level of fasting blood glucose.

**Supplemental Table 10.** Sensitivity analyses for intraindividual repeatability (intraclass correlation coefficient) in baseline and stress response glucose levels assessed between mid- and late development.

| | Baseline glucose | | Glucose response to handling stress (mg/dl) [a] | |
|---|---|---|---|---|
| | *N* | *ICC (95% CI)* | *N* | *ICC (95% CI)* |
| **Glucose measured in < 4min** [b] | 105 | 0.22 (0.0, 0.42) | 105 | 0.00 (-0.02, 0.00) |

[a] Glucose response to handling stress is calculated as the difference between the stress-induced level minus the baseline level of fasting blood glucose.

[b] Sensitivity analysis in which model is based on restricted data set that includes only baseline glucose measures collected in under 4 minutes.

**Supplemental Table 11.** Sensitivity analyses for models of the assocations between size and growth and fasting blood glucose levels assessed in nestling barn swallows during mid- and late development.

| | Mid-development (~8 days post hatch) | | | | Late development (~12 days post-hatch) | | | |
|---|---|---|---|---|---|---|---|---|
| | Baseline glucose (mg/dl) | | Glucose response to handling stress (mg/dl) [a] | | Baseline glucose (mg/dl) | | Glucose response to handling stress (mg/dl) [a] | |
| **Models** | ***N*** | ***β (95% CI)*** [b] | ***N*** | ***β (95% CI)*** [b] | ***N*** | ***β (95% CI)*** [b] | ***N*** | ***β (95% CI)*** [b] |
| **Mid-development right-wing length** | | | | | | | | |
| Adujsted for number nestlings [c] | 104 | **6.53 (0.85, 12.28)** | 100 | 0.10 (-5.48, 5.68) | 103 | **8.17 (2.45, 13.75)** | 103 | 4.45 (-1.49, 10.19) |
| Glucose measured in < 4min [d] | 85 | **6.34 (0.30, 12.26)** | 82 | 0.16 (-5.71, 6.03) | 93 | **7.05 (1.28, 12.78)** | 93 | **6.97 (0.88, 12.95)** |
| **Mid-development relative size** [e] | | | | | | | | |
| Adujsted for number nestlings [c] | 104 | 7.87 (-1.18, 17.19) | 100 | -3.75 (-13.50, 6.00) | 103 | 8.65 (-0.67, 17.95) | 103 | -1.44 (-13.07, 9.45) |
| Glucose measured in < 4min [d] | 85 | 10.52 (-0.05, 21.30) | 82 | -5.02 (-15.99, 5.96) | 93 | 7.90 (-1.60, 17.58) | 93 | 0.23 (-12.62, 11.28) |
| **Late development right-wing length** | | | | | | | | |
| Adujsted for number nestlings [c] | - | - | - | - | 106 | **11.37 (5.67, 17.33)** | 106 | **6.55 (0.03, 12.67)** |
| Glucose measured in < 4min [d] | - | - | - | - | 96 | **10.65 (4.88, 16.57)** | 96 | **8.32 (2.00, 14.53)** |
| **Late development relative size** [e] | | | | | | | | |
| Adujsted for number nestlings [c] | - | - | - | - | 106 | 4.31 (-4.61, 13.32) | 106 | 6.36 (-4.72, 16.78) |
| Glucose measured in < 4min [d] | - | - | - | - | 96 | 6.39 (-2.49, 15.49) | 96 | 8.49 (-1.99, 18.77) |
| **Growth based on right wing-length difference** | | | | | | | | |
| Adujsted for number nestlings [c] | - | - | - | - | 103 | **7.52 (0.84, 14.68)** | 103 | 4.04 (-2.80, 10.28) |
| Glucose measured in < 4min [d] | - | - | - | - | 93 | **8.38 (1.70, 15.22)** | 93 | 3.73 (-3.02, 10.39) |

[a] Glucose response to handling stress is calculated as the difference between the stressed induced concentration minus the baseline concentration of blood glucose levels.

[b] Estimated β (95% CI) from linear mixed models in which nest ID was included as a random intercept. Models include the age of the nest, as determined from the day that the first nestling hatched, and the time (s) elapsed between nestling removal from the nest and the blood draw.

[c] Sensitivity analysis in which model is adjusted for the number of nestlings in a nest.

[d] Sensitivity analysis in which model is based on restricted data set that includes only baseline glucose measures collected in under 4 minutes.

[e] Relative size is calculated based on an individuals right wing length below (small) or above (large) the average right wing length within a nest. Small is set as the reference category.

Bold estimates are significant at $p < 0.05$.

**Supplemental Table 12.** Associations between late development nestling size and growth with late development blood glucose assessed in models stratified by levels of parental care and *P* values for the prodcut term from an interaction model.

| | Low parental care models | | High parental care models | | Interaction model |
|---|---|---|---|---|---|
| **Glucose by size/growth models** | ***N*** | ***β (95% CI)*** [a] | ***N*** | ***β (95% CI)*** [a] | $P_{intx.}$ [d] |
| Baseline glucose by 1 SD right-wing length | 42 | **15.38 (6.25, 24.90)** | 42 | 4.70 (-3.17, 13.27) | 0.35 |
| Baseline glucose by 1 SD growth [b] | 42 | 8.43 (-0.18, 16.96) | 39 | -2.08 -14.25, 12.47) | 0.36 |
| Glucose response by 1 SD right-wing length | 42 | **12.53 (2.51, 22.31)** | 42 | 4.17 (-5.05, 14.45) | 0.55 |
| Glucose response by 1 SD growth [b, c] | 42 | **12.10 (2.53, 20.52)** | 39 | -2.81 (-15.65, 10.10) | 0.08 |

[a] Estimated β (95% CI) from linear mixed models of the association between size/growth glucose, stratifed by level of parental feeding. Nest ID was included as a random intercept. Models adjusted for the time at baseline blood draw and nest age.

[b] Growth is the difference in the right-wing length between mid- and late development.

[c] Glucose response to handling stress is calculated as the difference between the stressed induced concentration minus the baseline concentration of fasting blood glucose

[d] The product term from the interaction model is for right-wing length/growth by parrental care.

Bold estimates are significant at $P < 0.05$.
